# Integration of spatial and single-cell data across modalities with weak linkage

**DOI:** 10.1101/2023.01.12.523851

**Authors:** Shuxiao Chen, Bokai Zhu, Sijia Huang, John W. Hickey, Kevin Z. Lin, Michael Snyder, William J. Greenleaf, Garry P. Nolan, Nancy R. Zhang, Zongming Ma

**Author notes:** Senior Authors (Garry P. Nolan, Nancy R. Zhang, Zongming Ma). Equal Contributions (Shuxiao Chen, Bokai Zhu).

## Abstract

single-cell sequencing methods have enabled the profiling of multiple types of molecular readouts at cellular resolution, and recent developments in spatial barcoding, in situ hybridization, and in situ sequencing allow such molecular readouts to retain their spatial context. Since no technology can provide complete characterization across all layers of biological modalities within the same cell, there is pervasive need for computational cross-modal integration (also called diagonal integration) of single-cell and spatial omics data. For current methods, the feasibility of cross-modal integration relies on the existence of highly correlated, a priori “linked” features. When such linked features are few or uninformative, a scenario that we call “weak linkage”, existing methods fail. We developed MaxFuse, a cross-modal data integration method that, through iterative co-embedding, data smoothing, and cell matching, leverages all information in each modality to obtain high-quality integration. MaxFuse is modality-agnostic and, through comprehensive benchmarks on single-cell and spatial ground-truth multiome datasets, demonstrates high robustness and accuracy in the weak linkage scenario. A prototypical example of weak linkage is the integration of spatial proteomic data with single-cell sequencing data. On two example analyses of this type, we demonstrate how MaxFuse enables the spatial consolidation of proteomic, transcriptomic and epigenomic information at singlecell resolution on the same tissue section.

## Introduction

Recent technological advances have enabled the profiling of multiple biological modalities within individual cells, over many cells in parallel. The growing list of modalities that can now be profiled at the single-cell level include proteome and metabolome (1, 2), transcriptome (3), and various aspect of the epigenome such as methylation (4), histone modification (5–7), and chromatin accessibility (5, 8). In addition to technologies that operate on dissociated single cells, rapid progress has been made on the in situ measurement of transcriptome (9), proteome (10–14), epigenome (15), and other modalities on histological tissue sections at single-cell or close to single-cell resolution, retaining the spatial context. These advances have spawned consortia-level efforts to construct multiomic single-cell and spatial atlases of each and every organ, across species, in healthy and diseased states. To harness the new technologies and growing data resources for biological discovery, a primary challenge is the reliable integration of data across modalities. Cross-modal integration, also referred to as “diagonal integration” (16, 17), is the alignment of single cells or spatial spots across datasets where different features (or modalities) are profiled in each dataset. An example is the alignment of cells in a CODEX dataset, which measures protein abundance, to cells in a single-cell RNA sequencing (scRNA-seq) dataset, which measures RNA expression. This cross-modal integration step underpins many types of downstream analyses, and its importance is evident in the myriad methods that have already been developed to tackle it (18–24).

Despite the progress in this area, key limitations still hinder reliable cross-modal integration, as highlighted by recent surveys (16, 17, 25). A key factor limiting the accuracy of existing methods is the strength of *linkage* between modalities, as we define below. A feature is “linked” between two modalities if it can be measured in, or predicted by, both modalities. In the terminology of (16, 17), these linked features can serve as “anchors” for the integration. For example, to integrate single-cell or spatial ATAC sequencing (ATAC-seq) and single-cell or spatial RNA-seq data, most existing methods predict the “activity” for each gene in each cell/spot of the ATAC-seq data based on the accessibility of the gene’s surrounding chromatin; then, each gene’s ATAC activity can be linked to its RNA expression, mapping cells from the two datasets into the same feature space. Similarly, between RNA and protein assays, the abundance of each protein in the protein assay can be linked to the expression of its coding gene in the RNA assay. With the exception of bindSC (26), all existing methods, to our knowledge, rely crucially on the linked features and are designed for scenarios where there is a large number of linked features that exhibit strong cross-modality correlation, a situation that we refer to as “strong linkage”. For example, between scRNA-seq and scATAC-seq, every gene in the genome can be linked, and the correlation between gene activity and RNA expression is often high enough for enough genes to allow for precise integration (18, 19, 22). To achieve strong linkage, some methods attempt to learn a mapping from the features of one modality to the features of the other modality through a “training set” consisting of data where both modalities are simultaneously observed in each cell/spot (23, 27). While this strategy may be applicable towards the integration of data from biological systems that are similar to the training set, it is questionable how well it can generalize to unseen systems.

Cross-modality integration in scenarios of weak linkage, where the number of linked features is small and/or the between-modality correlation for the linked features is weak, is especially challenging. A prototypical example of weak linkage is between targeted protein assays (14, 28) and transcriptome/epigenome assays such as scRNA-seq and scATAC-seq. Such scenarios are becoming extremely common as spatial proteomic technologies are receiving wide-spread adoption (10–14), complementing RNA and ATAC sequencing in achieving more complete tissue characterization (see, for example, (29–32)). We will reveal, through comprehensive benchmarks, the limitations of existing state-of-the-art methods in such difficult cases.

Under both strong and weak linkage, the evaluation of existing methods have leaned heavily on systems with highly distinct cell types whose separation only requires a crude feature-level mapping between modalities. In fact, most existing methods explicitly focus on the goal of “label transfer”, that is, the transfer of cell type labels from one modality to the other. This goal only requires the integration to be accurate at the resolution of the label. As we demonstrate in our benchmarks, even this seemingly modest goal of label transfer for major cell types is unattainable in weak linkage scenarios by current methods, much less the more challenging goal of integration in continuously transitioning cell populations where subtle distinctions need to be preserved between closely related states. Yet, key biological discoveries often hinge on the accurate preservation of fine cell state distinctions during integration,

To address the above limitations, we developed MaxFuse (MAtching X-modality via FUzzy Smoothed Embedding), a model free, highly adaptive method that can accurately integrate data across weakly linked modalities. MaxFuse goes beyond label transfer and attempts to match cells to precise positions on a graph-smoothed low-dimensional embedding. MaxFuse starts by denoising the linked features in each modality through borrowing information from all of the features, and then performs an initial crude matching of cells based on the denoised linked features. Then, MaxFuse iteratively refines the matching step based on graph smoothing, linear assignment, and CCA. These iterations use information from all features in both modalities to improve upon the initial matching. The initial feature linkage may be derived from domain knowledge or an existing integration, and thus, MaxFuse can also be used to improve upon any existing integration methods.

We systematically benchmarked the performance of MaxFuse across protein, RNA, and chromatin accessibility single-cell multiome ground-truth datasets. Across a wide variety of datasets, MaxFuse has superior performance compared to other state-of-the-art integration methods. Although the largest improvements in accuracy are observed under weak linkage, under strong linkage MaxFuse is comparable to the current best method in integration performance with substantial improvement in speed.

We further demonstrate the analyses enabled by MaxFuse with two examples. First, in the integration of scRNA-seq and CODEX multiplexed in situ protein profiling data from the human tonsil, we show that MaxFuse can recover correct spatial gradients in the RNA expression of genes not included in the 46-marker protein panel. Next, MaxFuse is applied to an atlas-level integration of spatial proteomic and single-cell sequencing datasets, as part of a consortium-level effort to map cell organization and function across different regions of the human intestine (32). We demonstrate how to perform trimodal integration of CODEX, snRNA-seq, and snATAC-seq data to recover spatial patterns of RNA expression and transcription factor binding site accessibility at single-cell resolution.

## Results

### Cross-modality matching of single cells via iterative fuzzy smoothed embedding

Let data from the two modalities be represented by a pair of cell-by-feature matrices that contain all measured features in each modality. For convenience, call the two modalities *Y* and *Z*. In addition, we represent the initial knowledge about the linkage between the two modalities as another pair of cell-by-feature matrices whose columns have one-to-one correspondences. To distinguish between these two pairs of matrices, we call the former all-feature matrices and the latter linked-feature matrices. For example, when one modality is protein abundance over a small antibody panel and the other is RNA expression over the whole transcriptome, the two all-feature matrices have drastically different numbers of columns, one being the number of proteins in the panel and the other being the number of genes in the transcriptome; the linked feature matrices, on the other hand, have equal number of columns, where each column in the protein matrix is one protein and its corresponding column in the RNA linked-feature matrix is the gene that codes for the protein. When the number of cells is large, we recommend aggregating cells with similar features into meta-cells, as described in Materials & Methods, prior to applying MaxFuse. In that case, each row in the above matrices would represent a meta-cell. The procedure below does not depend on whether single- or meta-cells are used, and thus we will refer to each row as a “cell”. The two pairs of matrices form the input of the MaxFuse pipeline in Figure 1A.

**Figure 1:**
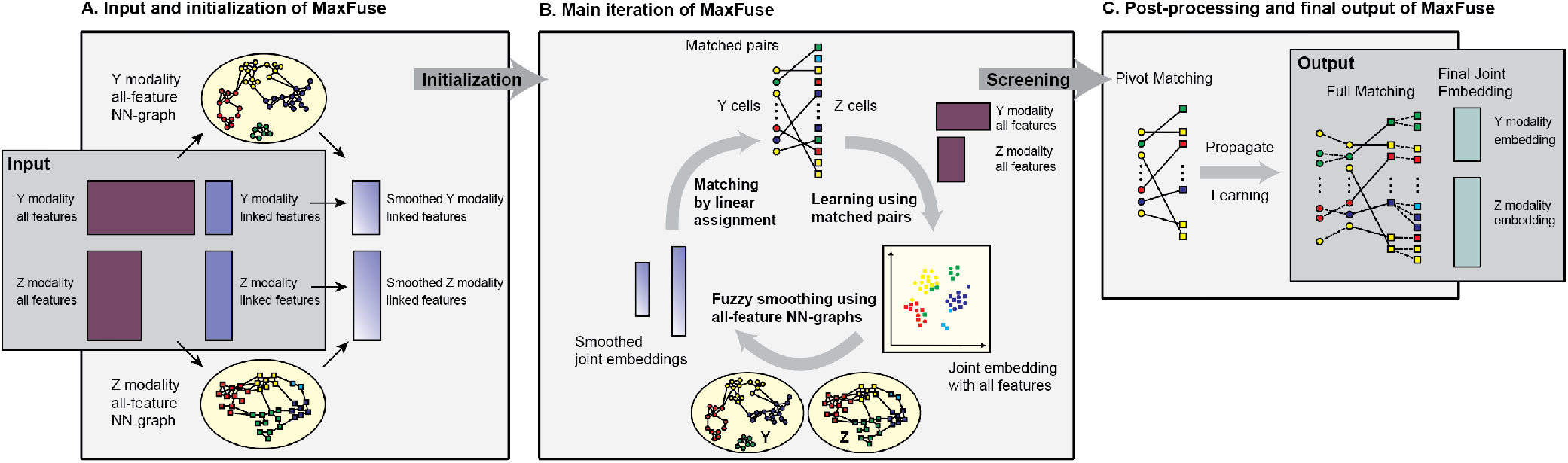
Overview of MaxFuse pipeline. **(A)** The input consists of two pairs of matrices. The first pair consists of all features from each modality, and the second pair consists of only the linked features. MaxFuse uses all features within each modality to create a nearest-neighbor graph (all-feature NN-graph) for cells in that modality. Fuzzy smoothing induced by the all-feature NN-graph is applied to the linked features in each modality. Cross-modal cell matching based on the smoothed linked features initializes the iterations in **(B). (B)** In each iteration, MaxFuse starts with a list of matched cell pairs. A cross-modal cell pair is called a pivot. MaxFuse learns CCA loadings over all features from both modalities based on these pivots. These CCA loadings allow the computation of CCA scores for each cell (including cells not in any pivot), which are used to obtain a joint embedding of all cells across both modalities. For each modality, the embedding coordinates then undergo fuzzy smoothing based on the modality-specific all-feature NN-graphs (obtained in **(A)**). The smoothed embedding coordinates are supplied to a linear assignment algorithm which produces an updated list of matched pairs to start the next iteration. **(C)** After iterations end, MaxFuse screens the final list of pivots to remove low-quality matches. The retained pairs are called refined pivots. Within each modality, any cell that is not part of a refined pivot is connected to its nearest neighbor that belongs to a refined pivot and is matched to the cell from the other modality in this pivot. This propagation step results in a full matching. MaxFuse further learns the final CCA loadings over all features from both modalities based on the refined pivots. The resulting CCA scores give the final joint embedding coordinates.

Stage 1 of MaxFuse aims to summarize cell-cell similarity within each modality and learn an initial cross-modal matching of cells. As shown in Figure 1A, this stage consists of three major steps. In step 1, for each modality, we use all features to compute a fuzzy nearest-neighbor graph connecting all cells measured in that modality. This graph, by utilizing the information in all features, provides the best possible summary of the cell-cell similarity for the given modality. In particular, cells that are close in this graph should have comparable values for their linked features. Thus, in step 2 of stage 1, MaxFuse boosts the signal-to-noise ratio in the linked features within each modality by shrinking their values, for each cell, towards the cell’s graph-neighborhood average. We call this step “fuzzy smoothing”. After fuzzy smoothing of linked features within each modality, MaxFuse computes in step 3 distances between all cross-modal cell pairs based on the smoothed linked features and applies linear assignment (33) on the cross-modal pairwise distances to obtain an initial matching of cells. The initial matching serves as the starting point of stage 2 of MaxFuse.

Stage 2 of MaxFuse, shown in Figure 1B, aims at improving cross-modal cell matching quality by iterating the sequence of joint embedding, fuzzy smoothing, and linear assignment steps. Starting with the initial matches obtained in stage 1, in each iteration, MaxFuse first learns a linear *joint embedding* of cells across modalities by computing a canonical correlation based on all features of the cross-modal matched cell pairs. Then, coordinates of this joint embedding are treated as new linked features of each modality and *fuzzy smoothing* is applied on them based on the all-feature nearest-neighbor graphs computed in stage 1. Finally, MaxFuse updates the cell-matching across modalities by applying *linear assignment* on the pairwise distances of these fuzzy-smoothed joint embedding coordinates. The resulting matching then starts the next iterate. Matching quality improves with each iteration until available information in all features, and not just the linked features, have been used.

Stage 3 of MaxFuse aims at post-processing the last crossmodal cell matching from stage 2 and producing final outputs. First, MaxFuse screens the matched pairs from the last iterate in stage 2, retaining high quality matches as pivots. The pivots are used in two complementary ways: (i) they are used one last time to compute a final joint embedding of all cells in both modalities; (ii) for any unmatched cell in either modality, its closest neighbor within the same modality that belongs to a pivot is identified and, as long as its distance to this neighbor is below a threshold, the match in the pivot is propagated to the cell. Thus, the final output of MaxFuse has two components: (i) a list of matched pairs across modalities, and (ii) a joint embedding of all cells in both modalities. More details on the MaxFuse algorithm are given in Materials & Methods.

### Integration of transcriptome and targeted protein data with varying protein panel sizes

We benchmarked Max-Fuse on a CITE-seq dataset (34) containing simultaneous measurements of 228 protein markers and whole transcriptome on peripheral blood mononuclear cells. For comparison, we also applied four state-of-the-art integration methods: Seurat (V3) (24), Liger (22), Harmony (20), and BindSC (26) to this same dataset. Protein names were converted to RNA names manually to link the features between datasets. In each repetition of our experiment, we randomly subsampled 10, 000 cells, applied all methods, and assessed using the benchmarking criteria to be described below. We performed 5 such repetitions and averaged the criteria across repetitions. We masked the known cell-cell matching between the protein and RNA modalities when applying all methods (treating Protein and RNA as two unpaired modalities), and then used the known matching for assessment.

Methods are assessed using six different criteria that measure both cell-type-level label transfer accuracy as well as cell-level matching accuracy. The first two criteria are based on label transfer accuracy. Cells are annotated at two levels of granularity: level-1, which differentiates between 8 major cell types, and level-2, a finer classification which differentiates between 20 cell types. Label transfer accuracy is expected to be higher for level-1 labels than for level-2 labels. The proportion of matched pairs that share the same label at both annotation levels are reported, with higher proportions indicating higher matching quality. The next two criteria measure the quality of the cross-modal joint embedding of cells. A high-quality joint embedding should preserve biological signal, as reflected by the separation of known cell types, while mixing the two modalities as uniformly as possible. Usually, there is a trade-off between biological signal preservation and uniformity of mixing. Thus, we report the *F*_1_ scores computed based on average silhouette width (slt_f1) and adjusted Rand index (ari_f1), as proposed in Tran et al. (35). These scores aggregate quality assessments of biological signal preservation and modality mixing. For both criteria, higher *F*_1_ indicates a better embedding. The fifth criterion is FOSCTTM, <u>F</u>raction <u>O</u>f <u>S</u>amples <u>C</u>loser <u>T</u>han <u>T</u>rue <u>M</u>atch (19, 36, 37), that quantifies the quality of joint embedding at single-cell resolution. For each cell, one can compute the fraction of cells in the other modality that is closer than its true match in the joint embedding. FOSCTTM is the average of this fraction over all cells in both modalities. The lower this measure, the closer the true matches are in the joint embedding, and hence, the better the joint embedding. The last criterion is FOSKNN, <u>F</u>raction <u>O</u>f <u>S</u>amples whose true matches are among their <u>*K*</u>-<u>N</u>earest <u>N</u>eighbors in the joint embedding space. For any given *k* ≥ 1, the higher this proportion, the better the joint embedding. For precise definitions and details of these criteria, see Materials & Methods. Among all criteria described above, MaxFuse uniformly dominates the methods by a sizable margin (Figure 2A). Importantly, MaxFuse provides accurate cell matching across weakly-linked modalities (level 1 accuracy 93.9%, + ∼ 7% to the second best method, Figure S2B). The UMAP plots calculated based on the post-integration embedding from respective methods are shown in Figure 2B, colored by modality and by cell type. MaxFuse achieves both better mixing of the two modalities (left panel) and better preservation of biological signals (right panel). For example, B cell subtypes (B naive, intermediate, and memory cells) present a nicely resolved developmental trajectory after MaxFuse integration, but not after integration by other methods.

**Figure 2:**
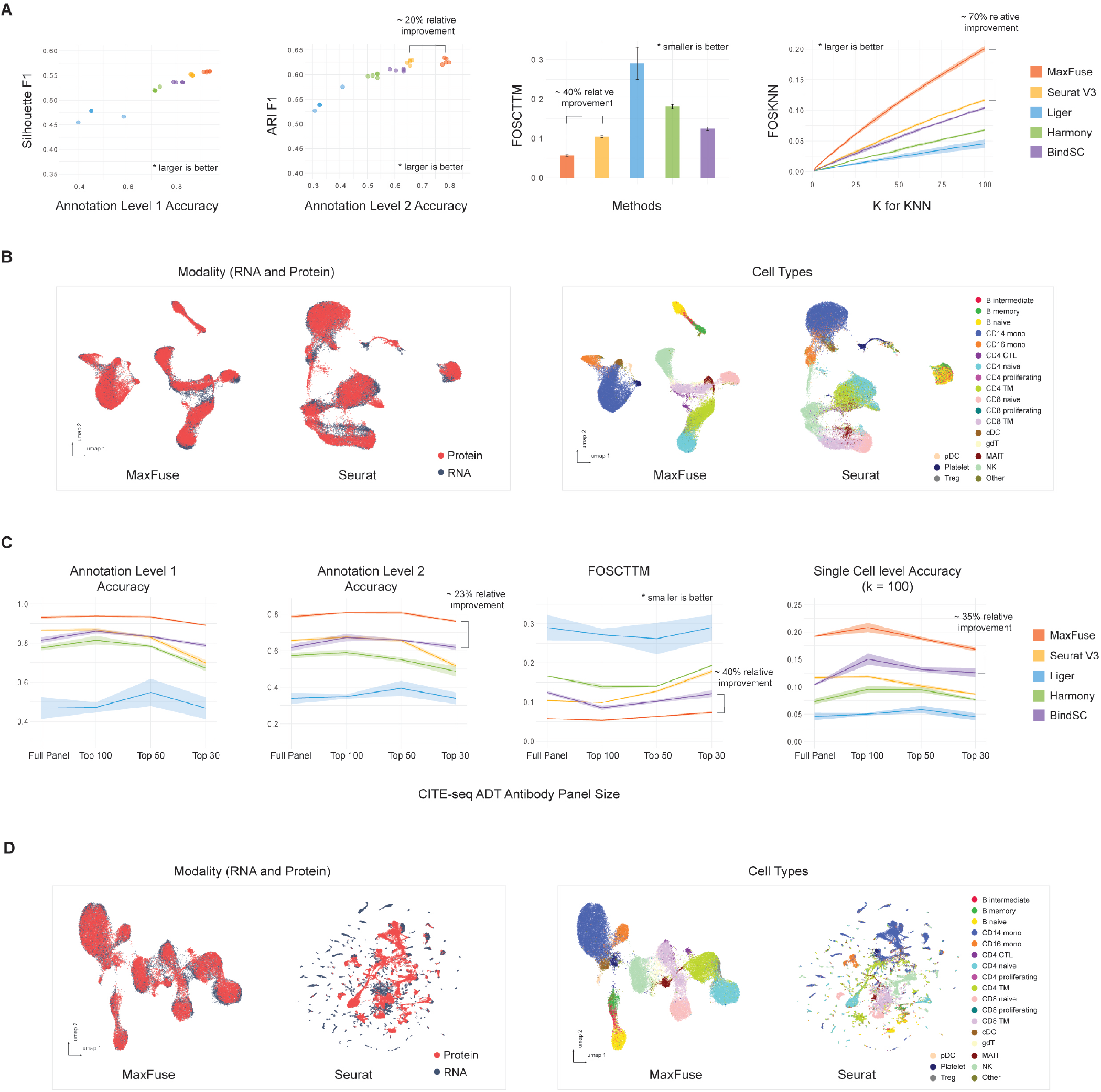
Benchmark on ground-truth CITE-seq PBMC data with full and reduced antibody panels. **(A)** Matching and integration performance of MaxFuse and other methods on CITE-seq PBMC dataset with full antibody panel (228 antibodies). **(B)** UMAP visualization of MaxFuse and Seurat (V3) integration results of CITE-seq PBMC with full panel, colored by modality (left) or cell type (right). **(C)** Matching and integration performance of MaxFuse and other methods on CITE-seq PBMC dataset with reduced antibody panels. **(D)** UMAP visualization of MaxFuse and Seurat (V3) integration results of CITE-seq PBMC with the 30 most informative of the original 228 antibodies, colored by modality (left) or cell type (right).

It is common to have an antibody panel that is of significantly smaller size than 228, especially for the emerging spatial-proteomic datasets. To benchmark the performance of MaxFuse against existing methods for smaller antibody panels, we ordered the proteins according to their importance for differentiating cell types (See Materials & Methods for details). We repeated the foregoing experiments when only the top 100, 50, and 30 most important proteins are used in the matching and integration process. At each antibody panel size, we ran the experiment over five independent repetitions with randomly subsampled 10, 000 cells, and average the cell type annotation level matching accuracy, FOS-CTTM and FOSKNN across repetitions (Figure 2C). Regardless of panel change, MaxFuse consistently outperformed other methods. Additionally, MaxFuse successfully mitigated the effect of reduced panel size on integration quality: Even when the antibody panel size was reduced to 30, MaxFuse maintained a *>* 90% annotation level 1 accuracy while other methods produced variable and low quality cell matching results (∼ 10 − 70%, Figure S2B). Similarly, with a reduced antibody panel size (eg. 30 antibodies), the integrated UMAP embedding (38) produced by other methods blurs the distinction between cell types, while MaxFuse embedding still accurately captures the subtle structure of highly granular cell subtypes (e.g., the B cell subpopulations, Figure 2D and Figure S2A).

### Systematic benchmark across multiple ground-truth multiome modalities

We further benchmarked MaxFuse on four additional single-cell multiome datasets. The first is a CITE-seq dataset of human bone marrow mononuclear cells (BMMCs) that provides cell-matched measurements of the full transcriptome along with an antibody panel of size 25 (34). The second is an ABseq dataset, also of BMMCs, with an antibody panel of size 97 and the whole transcriptome (39). The third is an ASAP-seq PBMC dataset (40) with 227 antibodies and the whole epigenome measured in ATAC fragments. The fourth is a TEA-seq PBMC dataset (41) where we focused on the simultaneous measurements of 46 antibodies and the whole epigenome measured in ATAC fragments. Together, these datasets represent a diverse collection of measurement technologies over different modality pairs. We benchmarked the performance of MaxFuse against Seurat (V3), Liger, Harmony, and BindSC on these datasets. For datasets with simultaneous RNA and protein features, we linked each protein to its coding gene. For datasets with simultaneous ATAC and protein measurements, we linked each protein to the gene activity score (42) computed from the ATAC fragments mapping near its coding gene. As in the previous case, the known cell-cell correspondence across modalities were masked in the matching and integration stage for all methods, but used afterwards for evaluation.

We compared the performances of MaxFuse and the other four methods on these datasets using the collection of matching and integration quality measures described in the previous section (Figure 3A): cell type annotation matching accuracy, FOSCTTM, FOSKNN (*K* set as 1*/*200 dataset size), Silhouette F1 score, and ARI F1. Overall, MaxFuse outperformed other methods, often by a sizable margin (eg. ∼ 20% relative improvement in terms of the metrics measured, Figure 3A and Figure S3.1A).

**Figure 3:**
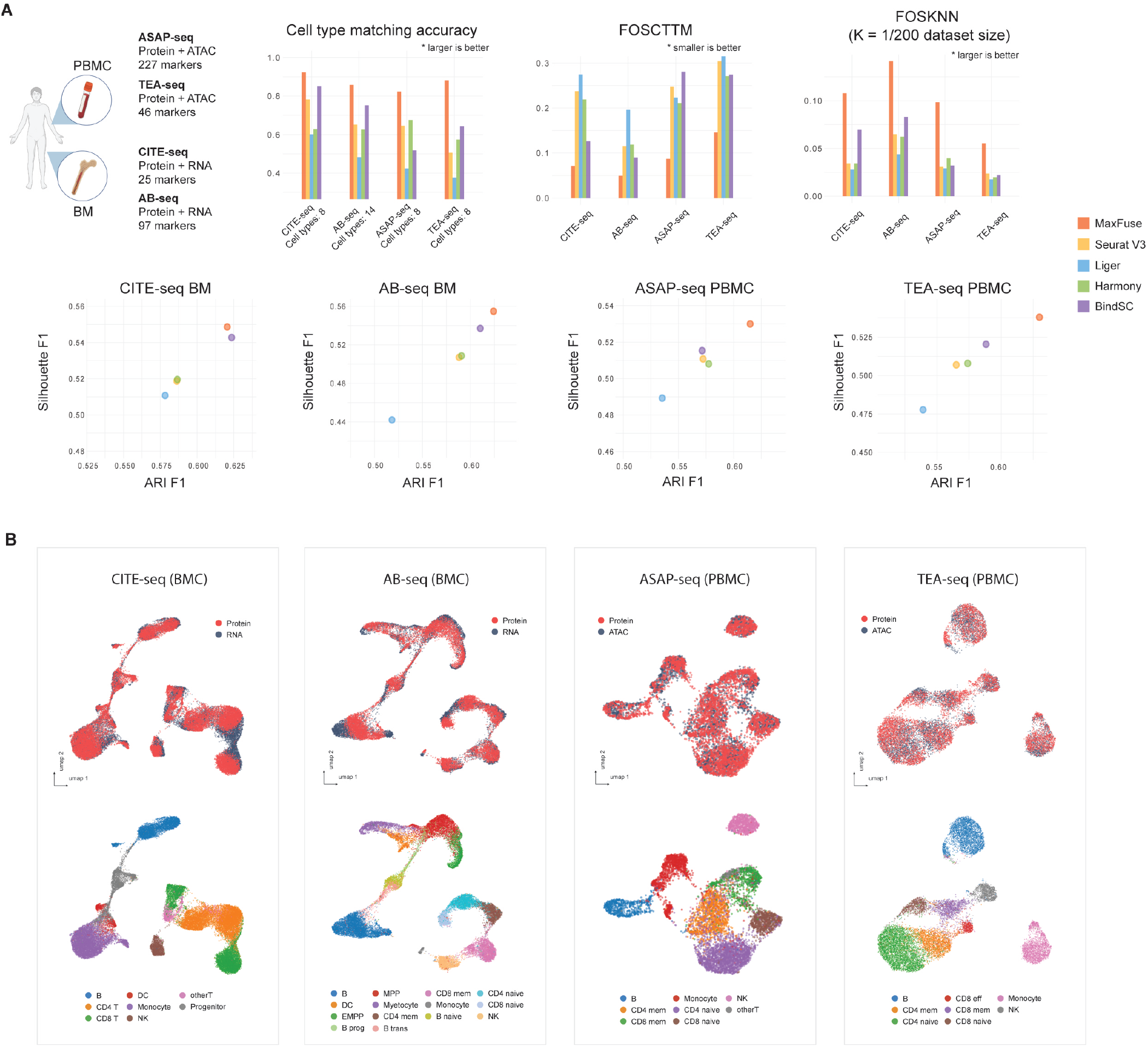
Systematic benchmark across multiple ground-truth data types with MaxFuse. **(A)** Four different multiome datasets, generated by different technologies, were benchmarked. Cell type matching accuracy, FOSCTTM, FOSKNN (with *K* = 0.5% total cell counts of each dataset), and ARI and Silhouette F1 were evaluated across 5 methods. **(B)** UMAP visualization of MaxFuse integration results for the four ground-truth multiome datasets.

UMAPs of the MaxFuse cross-modal joint embeddings for each dataset are shown in Figure 3B, with the top row colored by modality and the bottom row colored by cell type annotation. Across the integration scenarios, MaxFuse mixed different modalities well in joint embeddings while retaining separation between cell types. Compared to the UMAPs of joint embeddings produced by other methods, MaxFuse consistently achieves substantial improvements (Figure 3B and Figure S3.2 A).

As a counterpoint to the above integration scenarios, we also considered the problem of integration of scRNA-seq and scATAC-seq data, on which multiple methods have demonstrated feasibility (18, 19, 22). The degree of overlap in the information contained in the RNA and ATAC modalities has been systematically measured in Lin and Zhang (43), where it was shown that, in terms of cell population structure, the information shared across RNA and ATAC is much higher than the information shared between RNA and protein for commonly used targeted protein panels. Thus, RNA and ATAC has stronger linkage and should be easier to integrate. We benchmarked MaxFuse against state-of-the-art methods for this problem on four public multiome datasets that simultaneously measure the chromatin accessibility and transcriptome expression for each cell: 10x mononuclear cells from peripheral blood (44), cells from embryonic mouse brain at day 18 postconception (44), cells from developing human cerebral cortex (45), and cells from human retina (46). The integration quality criteria described in the previous subsection are used to assess all methods, shown in Supplementary Materials. Across datasets and evaluation metrics, MaxFuse achieves best or close-to-best performance among methods, and is comparable to scGLUE. However, MaxFuse is much faster than scGLUE. For example, for the integration of a dataset of 20,000 cells, MaxFuse took <5 minutes to finish on a laptop with M1 Max chip while scGLUE took hours on a comparable platform without CUDA acceleration.

### Cross-modal integration of scRNA-seq and spatial proteomic data enables information-rich spatial pattern discovery

MaxFuse is particularly motivated by scenarios where the signal-to-noise ratio in the cross-modal linked features is low. Weak linkage is especially common in spatial-omic data types due to technical limitations. For example, high resolution spatial proteomic methods such as CODEX, MIBI-TOF, IMC, and CosMx SMI can profile, at sub-cellular resolution, a panel of 30-100 proteins (10–13). Integration of such spatial proteomics datasets with single-cell transcriptomic and epigenomic datasets of the same tissue is often of interest, and particularly challenging due to the small number of markers in the spatial dataset and the weak linkage between modalities that is caused by both biological and technical differences. Thus, we demonstrated and benchmarked MaxFuse on the integration of CODEX multiplex imaging with 46 markers (47) with single-cell RNA-seq (48) of human tonsils from two separate studies (Figure 4A). Figure 4B shows the UMAPs of the MaxFuse integration colored by modality and by 6 major cell types.

**Figure 4:**
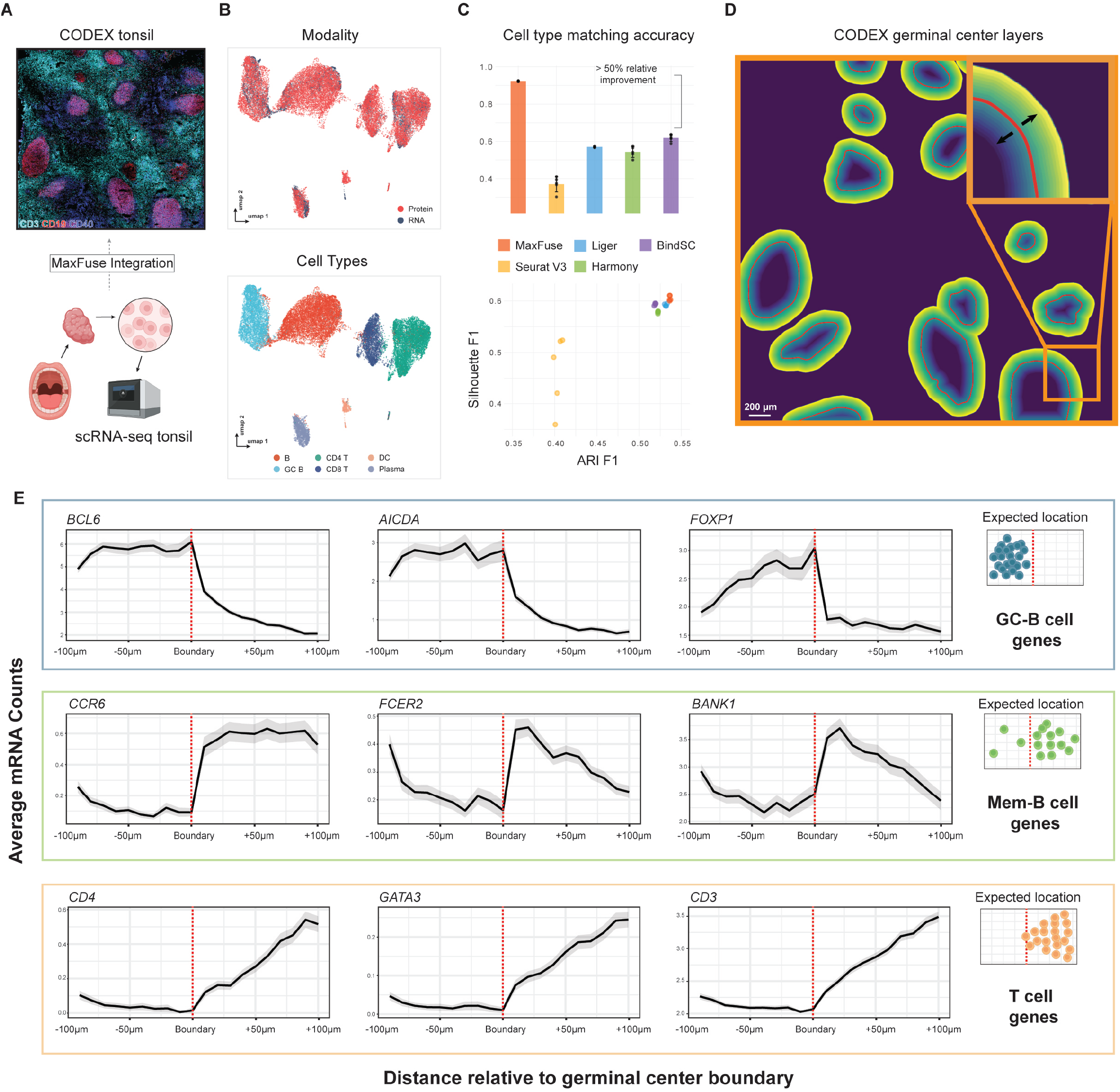
MaxFuse enables information-rich spatial pattern discovery. **(A)** MaxFuse integrates human tonsil single-cell data: one dataset by CODEX from Kennedy-Darling et al (47) (upper panel), the other dataset by scRNA-seq from King et al (48) (lower panel). **(B)** UMAP visualization of MaxFuse integration of tonsil CODEX and scRNA-seq data, colored by modality (upper panel) and cell type (lower panel). **(C)** Metrics (cell type matching accuracy, Silhoutte F1 and ARI F1 score) evaluating performance of MaxFuse and other methods. Five batches of randomly sampled CODEX and scRNA-seq cells (total of 40k each batch) were sampled, and used for benchmarking for all methods. **(D)** Illustration of cell layers extending inwards/outwards from the germinal center boundary, with each layer consisting of 30 pixels (*∼* 11*μ*m). A total of 10 layers extending in each direction were examined. **(E)** For each of 9 genes, the average mRNA counts (linked by MaxFuse) across cells in each layer are plotted versus the position of the layer in reference to the germinal center boundary (inward on the left of boundary, outward on the right). For each group of 3 genes (row), their expected expression profile in reference to the germinal center boundary is shown on the right.

Based on the pre-described benchmarking metrics, MaxFuse is the only method capable of integrating spatial proteomic and single-cell RNA-seq data. Existing state-of-the-art methods, Seurat (V3), Liger, Bindsc, and Harmony, failed to produce an embedding that integrates the two modalities while preserving the cell population structure (Figure 4B and Figure S4.1A). Evaluation results based on cell-type matching accuracy is consistent with evaluation results based on the joint embedding. At the level of the 6 major cell types shown in Figure 4B, MaxFuse is able to achieve high label transfer accuracy (93.3%), while the other methods fail to preserve cell type distinctions (40% - 60%, Figure 4B and Figure S4.1B).

We further assessed whether MaxFuse can preserve, during integration, the more subtle spatial variations within a cell type that are captured by CODEX. We manually delineated the boundaries of each individual germinal center (GC) from the CODEX tonsil images based on CD19, CD21, Ki-67 protein expression patterns. From the boundaries, we then extended outward or inward, with each step covering roughly one layer of cells (one step = 30 pixels erosion/dilation) (Figure 4C). Then, for each layer of cells, we calculated the average counts of specific genes, based on the scRNA-seq cells that match to CODEX cells of that layer. We then asked if known position-specific gene expression patterns relative to the germinal center boundary are recovered in the integrated scRNA-seq data. Indeed, MaxFuse was able to reconstruct the spatial pattern of the GC from disassociated transcriptomic data (Figure 4D): For GC-specific genes *BCL6, AICDA* and *FOXP1* (49–51) that relate to germinal center functionality, we observed high expression within the boundary and a sharp drop in expression after passing the boundary layer; for genes related to B cell memory *CCR6, BANK1* and *FCER2* (51–53) that should be enriched in B cells exiting from the GC, we indeed saw a gradual increase outside of the GC and then a quick decrease as the layer fully expands into the T cell region; and finally for T cell related genes, for example *CD4, GATA3 and CD3* (54), we indeed saw a rapid increase outside of the GC boundary but no expression within. In comparison, the integration with scRNA-seq produced by other methods was incapable of accurately reconstructing the GC spatial pattern (Figure S4.2A).

### Tri-modal atlas-level integration of spatial and single– cell data with MaxFuse

In the consortium-level effort to generate a comprehensive atlas across different regions of the human intestine, colon and small bowel tissue from healthy human donors were collected and systematically profiled by CODEX, snRNA-seq, and snATAC-seq (32). We applied MaxFuse to the integration of these three modalities (Figure 5A), with the goal of constructing high-resolution spatial maps of full transcriptome RNA expression and transcriptome factor binding accessibility. To perform tri-omic integration, we first conducted pairwise alignment of cells between protein (CODEX) and RNA (snRNA-seq), and cells between RNA (snRNA-seq) and ATAC (snATAC-seq), as previously described. The two sets of bi-modal cell-pairing pivots were then “chained” together, with the pivot cells in the RNA modality serving as the intermediary. This “chaining” created a set of pivots linking all three modalities: Protein, RNA, and ATAC. Subsequently, we used these pivots to calculate a tri-omic embedding via generalized CCA (gcca) (21, 55). This allows for a joint UMAP embedding of the three modalities, shown in Figure 5B. We see that distinctions between major cell types are preserved and modalities are mixed within each cell type.

**Figure 5:**
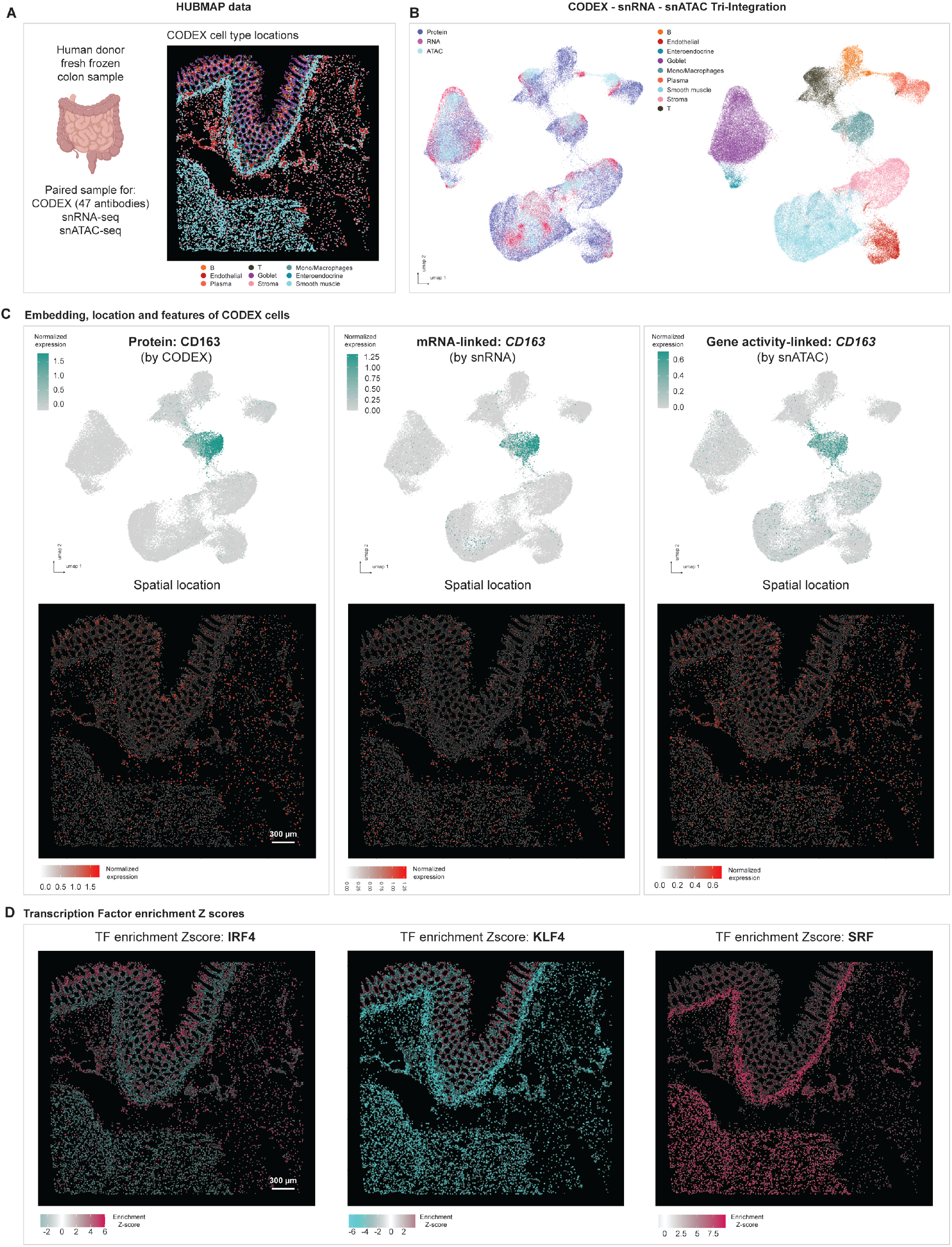
MaxFuse enables tri-modal integration with HUBMAP data. **(A)** Overview of data: Patient paired CODEX, snRNA-seq, snATAC-seq single-cell human intestine data from HUBMAP consortium. Colon and small bowel data were integrated by MaxFuse respectively and this figure shows part of the colon data (CODEX data from one donor; snRNA-seq and snATAC-seq data from four donors). **(B)** UMAP visualization of the tri-modal integration embedding produced by MaxFuse, colored by modality: Protein, RNA and ATAC (left panel) and colored by cell type (right panel). **(C)** Upper panel: UMAP visualization of CODEX cells based on the integration embedding, overlaid with CD163 protein expression (from CODEX cells itself, left panel), *CD163* RNA expression (from matched snRNA-seq cells, middle panel), *CD163* gene activity score (from matched snATAC-seq cells). Lower panel: Spatial location of CODEX cells based on their centroids’ x-y position, overlaid with the same expression features as in the upper panel. **(D)** Spatial location of CODEX cells based on their centroids’ x-y position, overlaid with the transcription factor motif enrichment score (Z-score, calculated by `chromVAR` (56)), based on their matched snATAC-seq cells.

The MaxFuse integration produces, effectively, a joint profile of protein abundance, RNA expression, and chromatin accessibility at single-cell spatial resolution on the same tissue section. To confirm the post-integration consistency between the three modalities, we inspected whether CODEX’s protein abundance aligns spatially with the expression and chromatin activity of the protein-coding gene, the spatial measurements of the latter two modalities imputed based on the MaxFuse integration. Figure 5C shows an example in CD163, a macrophage marker: The protein expression, RNA expression, and gene activity of CD163 are, as expected, uniquely enriched in the macrophage cell cluster (Figure 5C upper panel). Furthermore, protein, RNA, and ATAC activities of this gene all localize to the same spatial positions on the tissue section (Figure 5C lower panel). Other examples are shown in Supplementary Materials.

With the integration of the snATAC-seq and CODEX data, we can further map the spatial enrichment of transcription factor (TF) binding site accessibility. For each TF, this is achieved by first computing its motif enrichment score for each cell in the snATAC-seq data, and then the scores are transferred to the CODEX spatial positions based on the MaxFuse integration. Figure 5D shows such spatial profiles for 3 transcription factors: Binding motifs of *IRF4*, known to be a key regulator in immune cell differentiation (57), had increased accessibility in the immune-enriched compartments of the mucosa and submucosa layers (32). Binding motifs of *KLF4*, known to be required for the terminal differentiation of goblet cells (58), had heightened accessibility in the colonic crypts of the mucosa layer where goblet cells mature. Finally, binding motifs of *SRF*, a master regulator of smooth muscle gene expression, (59), had heightened accessibility in neighborhoods that are enriched for smooth muscle cells.

## Discussion

In this paper, we conceptually separated cross-modal integration of single-cell data into two different scenarios: across modalities with strong linkage (e.g., ATAC-RNA integration) and across modalities with weak linkage (e.g., RNA-protein integration for a targeted protein panel). Most existing methods are developed for integration across strongly linked modalities, and our ground-truth benchmark results suggest that their performances decay significantly as the strength of cross-modal linkage weakens. MaxFuse is motivated by and focuses on the challenging case of weak linkage, which has become increasingly common as many emerging study designs include spatial data with targeted marker panels to be collected jointly with single-cell sequencing data. MaxFuse relies on two key ideas to overcome weak linkage: The first is a “fuzzy smoothing” procedure that denoises the linked features by moving their values towards their graph-smoothed values, with the graph determined by all features. The second is an iterative refinement procedure that improves the cross-modal matching through an iterative cycle of coembedding, graph-smoothing, and matching; this ensures that information from *all* features, in both modalities, are used to generate the final matching. We show that these key ideas allow MaxFuse to substantially improve upon state-of-the-art methods, achieving accurate integration of data from targeted protein assays with data from transcriptome-and epigenome-level assays.

While MaxFuse is motivated by the weak linkage scenario, its applicability is universal. For strong linkage scenarios, methods based on deep learning, such as scGLUE, achieve state-of-the-art integration performance but is hindered by high computational costs. In comparison, MaxFuse achieves comparable performances as scGLUE on groundtruth strong-linkage benchmark datasets at a considerably lower computational cost. In addition, when joint embedding coordinates from other integration methods are available, these coordinates could serve as linked features in MaxFuse, which could then be further improved by the procedure. The light computation architecture and the flexibility in incorporating domain knowledge and existing integration results make the MaxFuse framework applicable to a wide range of cross-modal integration tasks.

## Materials & Methods

### The MaxFuse pipeline

#### Input preparation

Consider a pair of datasets 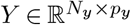 and 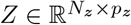 from two modalities (termed *Y* -modality and *Z*-modality for exposition convenience), with each row corresponding to a cell and each column a feature. In the ensuing discussion, we treat *Y* as the modality with a higher signal-to-noise ratio. For concreteness, one can think of *Y* as a snRNAseq dataset and *Z* as a CODEX dataset. Suppose there are two known functions 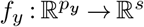 and 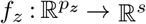 such that *f*_*y*_(*y*) predicts the values of *f*_*z*_(*z*) in a cell if the measured values under *Y* -modality are *y* in that cell and those under *Z*-modality are *z*. For any matrix *A* with *p*_*y*_ columns, let *f*_*y*_(*A*) denote the matrix with *s* columns and the same number of rows as *A*, obtained from applying *f*_*y*_ on each row of *A* and stacking the outputs as row vectors. For any matrix *B* with *p*_*z*_ columns, *f*_*z*_(*B*) is analogously defined. With *f*_*y*_ and *f*_*z*_, we define 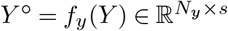 and 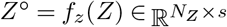. In the snRNAseq vs. CODEX example, if one has a crude prediction for a subset *S* (with size |*S*| = *s*) of the proteins then *f*_*z*_(*z*) = *z*_*S*_ returns the subvector indexed by *S* while 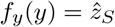 predicts the observed CODEX values for these proteins based on transcriptomic information of a cell. In summary, we start with a pair of original datasets (*Y, Z*) and a pair of datasets (*Y* °, *Z*°) with correspondence of columns based on domain knowledge.

##### Meta-cell construction

To alleviate sparsity and to scale to large datasets, we start by constructing meta-cells. Take the *Y* -modality for example. Let *n*_*y*_ be the desired number of meta-cells one aims for. We first construct a nearest-neighbor graph of the rows of *Y*, apply Leiden clustering with an appropriate resolution level to obtain *n*_*y*_ clusters, and average over the rows within each cluster to obtain the features for each meta-cell that serves as the representative of the cluster. Consequently, we obtain 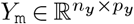. Using this clustering structure (induced by *Y* as opposed to *Y* °), we can average feature vectors in *Y* ° to obtain 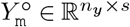. When desired, the same operation can be performed on the *Z*-modality to obtain 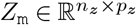 and 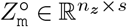. We recommend only constructing meta-cells for modalities with high signal-to-ratios. For example, if *Y* -modality contains snRNAseq data and *Z*-modality contains CODEX data, then we would construct meta-cells only in *Y* -modality. After this curation step, we have two pairs of datasets (*Y*_m_, *Z*_m_) and 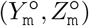. The former pair can have completely distinct feature sets, while the latter pair must have matching feature sets with corresponding columns. In Figure 1A, the former correspond to the pair of all feature matrices, and the latter correspond to the pair of linked feature matrices.

#### Fuzzy smoothing

Let 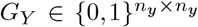 be a nearest neighbor graph of *Y*_m_ where each row *i* is connected to 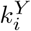 rows that are closest in a chosen similarity measure, including itself. So row *i* of *G*_*Y*_ has 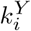 entries equal to one and others zeros. In addition, all its diagonal entries are equal to one. Let 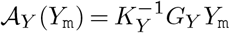 and 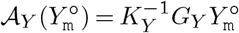 be locally averaged versions of *Y*_m_ and 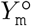 over *G*_*Y*_, respec-tively, where 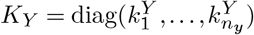. For a nearest neighbor graph *G*_*Z*_, we define A_*Z*_(*Z*_m_) and 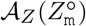 in an analogous way. Finally, for any weight *w* ∈ [0, 1) and any matrices *A* and *B* with *n*_*y*_ and *n*_*z*_ rows respectively, define

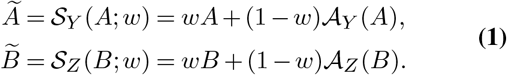

In this way, we define 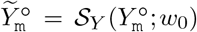 and 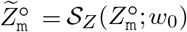 with *w*_0_ ∈ [0, 1). In Figure 1A, these are the smoothed *Y* -modality linked features and smoothed *Z*-modality linked features.

### Initial matching via linear assignment

As the columns in 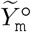 and in 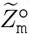 have correspondences, we can compute an *n*_*y*_ ×*n*_*z*_ distance matrix *D*° where 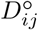 measures the distance between the *i*-th row in 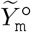 and the *j*-th row in 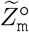 after projecting to respective leading singular subspaces. We obtain an initial matching 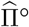 as the solution to the linear assignment problem (33, 60):

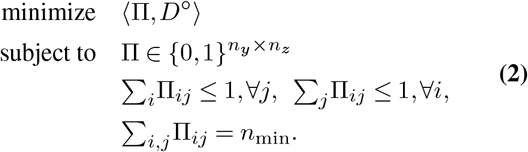

Here, *n*_min_ = min *n*_*y*_, *n*_*z*_ and for two matrices *A* and *B* of the same size, ⟨*A, B*⟩ = ∑ *i,j A*_*ij*_*B*_*ij*_ denotes the trace inner product. The estimator 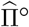 provides a relatively crude matching using only the information provided by the prior knowledge encapsulated in *f*_*y*_ and *f*_*z*_ that link features in the two modalities. By definition, 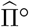 gives *n*_min_ pairs of matched rows between the two modalities. We call these matched pairs *initial pivots*.

#### Cross-modality joint embedding and iterative refinement of matching

##### From matched pairs to joint embedding

An estimated matching 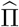 induces a cross-modality joint embedding of *Y*_m_ and *Z*_m_. In particular, let 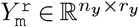 and 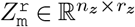 collect the leading PCs of *all* features (i.e., *Y*_m_ and *Z*_m_) in the two modalities, respectively. Here, the numbers of PCs to retain, i.e., *r*_*y*_ and *r*_*z*_, are chosen based on data. For any matrix *A*, let [*A*]_*i·*_ denote its *i*-th row. Suppose 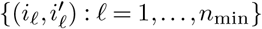 are the matched pairs specified by 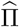. We perform CCA on data pairs

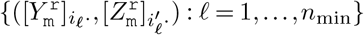

to obtain the leading *r*_cc_ loading vectors for either modality, collected as the columns of 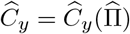 and 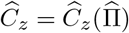, respectively. The cross-modal joint embedding induced by 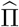 is then 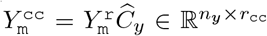 and 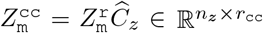, which are the predicted CC scores of 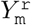 and 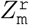 respectively.

##### Iterative refinement

Let 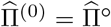 be the initial matching obtained from Eq. (2). Fix a weight *w*_1_ ∈ [0, 1) and the embedding dimension *r*^cc^, we refine the estimated matching by iterating the following steps for *t* = 1, …, *T* :

i. Compute joint embedding 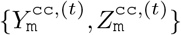 induced by 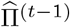;
ii. Apply fuzzy smoothing on joint embedding: 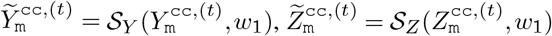;
iii. Calculate a distance matrix 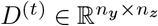 where 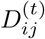 measures the distance between 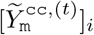. and 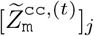., and obtain a refined matching 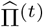 by solving Eq. (2) in which *D*° is replaced with *D*^(*t*)^.

Figure 1B illustrates the foregoing refinement iteration.

#### Propagation of matching and post-processing

For downstream analyses, one would often like to find for each cell in *Y* a match in *Z* when possible, or *vice versa*, and sometimes both ways. In addition, we would like to have joint embedding of cells across different modalities in a common space. We now describe how MaxFuse achieves these goals.

##### Filtering and final joint embedding

Upon obtaining the matched pairs 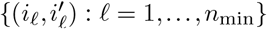 in 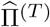, we rank them in descending order of 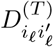 and only retain the top 100 × (1 − *α*)% pairs, where *α* is a user-specified filtering proportion (with a default *α* = 0). The retained pairs are called *refined pivots*. Then, we fit a CCA using the refined pivots and the corresponding rows in *Y*_m_ and *Z*_m_ to get the associated CCA loading matrices 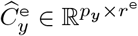 and 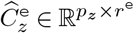. Here the positive integer *r*^e^ is a user-specified dimension for final joint embedding. Finally, the joint embedding of the full datasets is given by 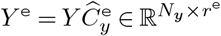 and 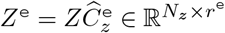, respectively. In Figure 1C, they correspond to the *Y* -modality embedding and *Z*-modality embedding matrices.

##### Using pivots to propagate matching

For each row index *i* ∈ {1, …, *n*_*y*_} in *Y* -modality that does not have a match in *Z*-modality (i.e., *i* does not belong to any refined pivot), we search for the nearest neighbor of the *i*-th row in 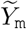 (*Y*_m_ after fuzzy smoothing) that belongs to some refined pivot. Suppose the nearest neighbor is the *j*_*i*_-th row with a match 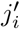 in *Z*-modality, then we call (*i*, 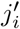) a matched pair obtained via *propagation*. We can optionally filter out any matched pair via propagation in which the nearest neighbor distance between 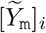 and 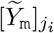 is above a user-specified threshold. The retained matched pairs composes the *Y* -to-*Z propagated matching*. We then repeat the above procedure with the roles of *Y* - and *Z*-modalities switched and obtain the *Z*-to-*Y* propagated matching.

Pooling all matched pairs from refined pivots and propagated matching together, we obtain a matching between meta-cells in *Y* -modality and those in *Z*-modality. Such a meta-cell level matching defines a single-cell level matching between the original datasets *Y* and *Z* by declaring (*i, i*′) a matched pair for 1 ≤ *i* ≤ *N*_*y*_, 1 ≤ *i*′≤ *N*_*z*_ if the meta-cell that *i* belongs to is matched to the meta-cell that *i*′ belongs to.

##### Scoring and directional pruning of matching

For each single-cell level matched pair (*i, i*′), we compute Pearson correlation between the *i*-th row of *Y* ^e^ and the *i*′-th row of *Z*^e^ (i.e., corresponding rows in final joint embedding) as its matching score. We use these matching scores to prune single-cell level matching, with the *direction* of pruning specified by user. Suppose the user wants to find for each cell in *Z* a match in *Y* (e.g., *Z* is a CODEX dataset and *Y* snRNAseq). Then for each cell index 1 ≤ *i*′≤ *N*_*z*_, we first list all refined pivots and propagated matching pairs that contain *i*′. If the list is non-empty, we only retain the pair with the highest matching score. Otherwise, we declare no match for cell *i*′ in *Z*-modality. If the direction is reversed, we apply the foregoing procedure with the roles of *Y* and *Z* switched. Furthermore, if no directional pruning is desired, we just keep all refined pivots and post-screening propagated matching pairs in the final single-cell matching.

After filtering, propagation, and potential pruning, the final list of matched pairs correspond to the final matching in Figure 1C.

### A batched version of MaxFuse

Single-cell and spatial datasets can be large. To facilitate fast computation for large datasets, we developed a batched version of MaxFuse.

#### Batching

Fix a desired pair of sample sizes (*n*_*y*_, *n*_*z*_) and meta-cell ratios (*N*_*y*_*/n*_*y*_, *N*_*z*_*/n*_*z*_), we randomly partition the dataset under *Y* -modality (resp. *Z*-modality) into disjoint subsets of sizes roughly all equal to *N*_*y*_ (resp. *N*_*z*_). Denote them as *Y* ^[1]^, …, 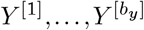 and *Z*^[1]^, …, 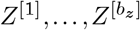. We then apply the MaxFuse pipeline on each pair of data {*Y* ^[*l*]^, *Z*^[*m*]^ }, 1 ≤ *l* ≤ *b*_*y*_, 1 ≤ *m* ≤ *b*_*z*_ to get the refined pivots and the propagated matching, as well as their induced single-cell level matched pairs, for that pair of batches.

#### Stitching

After pooling all refined pivots from all batch pairs, we obtain a multiple-to-multiple matching. For each unique cell in *Z*-modality, we average all its matches in *Y* - modality, that is, we average matched cells in the modality with a higher SNR. After this step, we get a pair of matrices with rows paired. We then fit CCA on this pair of matrices and get the loading matrices, which are then used to jointly embed the whole datasets. Finally, with the joint embedding of the whole datasets in *Y* - and *Z*-modalities, scoring and directional pruning of matching are performed in the same way as in MaxFuse without batching.

### Systematic benchmarks on ground-truth datasets

#### MaxFuse and other methods in comparison

MaxFuse was implemented in Python, and the four methods in comparison, Seurat V3, Harmony, Liger, and BindSC, were implemented in R. All benchmarking datasets were preprocessed in the same way for all methods, including filtering of low-quality cells, selection of highly variable genes and protein features to be used in integration, feature linkage scheme (e.g., protein to their corresponding gene names), and normalization of raw observed values (except for Liger which required scaling without centering). We used the default tuning parameters in each method suggested by the respective tutorial except for BindSC, for which we used the separate set of parameters suggested for the integration of protein-related data by its method tutorial website. For MaxFuse, initial matching used features that are weakly linked (e.g., protein CD4 and RNA *CD4*) and are smoothed by all-feature nearest-neighbor graphs. For refined matching, all features from both modalities were used (e.g., all proteins and RNAs that are highly variable). For other methods in comparison, BindSC used both the weakly linked features and all features, whereas others only used the weakly linked features by design. The full detail (including preprocessing, implementation, and downstream analysis and evaluation of MaxFuse and other methods) is recorded and can be reproduced.

#### Evaluation metrics

1. Cell type matching accuracy: To evaluate the matching performance for Seurat, Liger, Harmony, and BindSC, we used the respective integration embedding vectors produced by each method. For these methods, for each cell in one modality, we regarded its nearest neighbor from the other modality under Pearson correlation distance in the embedding space as its match. For MaxFuse, we directly used matched pairs produced in the final result. For all methods, we use the same matching direction (e.g., for each cell in CODEX data finding a matched cell in scRNAseq data) for fair comparison. Accuracy of the matchings was measured by fraction of matched pairs with identical cell type annotations. Details on cell type annotation are given below in the description of each benchmarking dataset.
2. FOSCTTM: Fraction of sample closer than true match (FOSCTTM) was used to evaluate single-cell level alignment accuracy on datasets with ground-truth single-cell level pairing. The measure has been used previously in cross-modality alignment benchmarking tasks (19, 36, 37). For such data, *N*_*y*_ = *N*_*z*_ = *N*, and FOSCTTM is defined as:

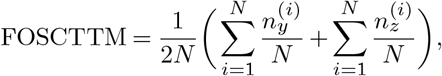

where for each *i*, 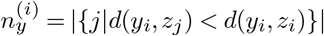 with *d* a distance metric in the joint embedding space and for *l* = 1, …, *N, y*_*l*_ and *z*_*l*_ the embedded vectors of the *l*-th cell with its measurements in *Y* and *Z* modality, respectively. The counts 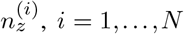, *i* = 1, …, *N*, are defined analogously. A lower value of FOSCTTM indicates better integration performance.
3. FOSKNN: Fraction of sample with true match among *k*-nearest-neighbors (FOSKNN) was used to evaluate single-cell level alignment accuracy on datasets with ground-truth single-cell level pairing. For such data, *N*_*y*_ = *N*_*z*_ = *N*. For any method in comparison, let {*y*_*i*_ : *i* = 1, …, *N*} be the coordinates of cells in the joint embedding space from their *Y* modality information, and let {*z*_*i*_ : *i* = 1, …, *N*} be embedding coordiantes from their *Z* modality information. Then

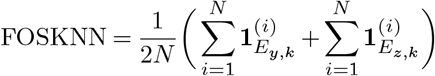

where for *i* = 1, …, *N*, 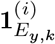 is the indicator of whether the *k* closest embedded vectors from *Z* modality to *y*_*i*_ includes *z*_*i*_. The quantity 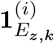 is defined analogously.
4. Silhouette F1 score: Silhouette F1 score has been used to simultaneously measure modality mixing and information preservation post-integration process (21, 35). In brief, the F1 score was calculated by 2 slt_mix slt_clust*/*(slt_mix + slt_clust), where `slt_mix` is defined as one minus normalized Silhouette width with the label being modality index (two modalities); `slt_clust` is defined by the normalized Silhouette width with label being cell type annotations (e.g., “CD4 T”, “CD8 T”, “B”, etc.). All Silhouette widths were computed using the `silhouette()` function from R package `cluster`.
5. ARI F1 score: Adjusted Random Index F1 score has been used to jointly measure modality mixing and information preservation post-integration process (21, 35). The score was calculated in a similar way to Silhouette F1 score, while the Adjusted Random Index was used instead of the Silhouette width. All ARI scores were computed using the function `adjustedRandIndex()` in R package `mclust`.

#### CITE-seq PBMC dataset

The CITE-seq healthy human pbmc data with antibody panel of 228 markers was retrieved from Hao et al. (34). For benchmarking purposes, 5 batches of cells, each with 10k cells were randomly sampled from the original dataset, and selected for benchmarking. The first 15 components of the embedding vector produced by all methods were used for benchmarking metric calculation. The UMAP visualization of the integration process was also calculated with the first 15 components of the embedding vectors. Cell type annotations (lv1 - 8 cell types and lv2 - 20 cell types) were directly retrieved from Hao et al.’s original annotation.

For antibody dropping, we ranked the importance of each individual antibody in the panel in terms of phenotyping contribution. The importance score was calculated by training a random forest model (function `randomForest` in R package `randomForest`, with default parameters) using all antibodies to predict cell type labels (annotation level 2 from Hao et al.), then a permutation feature importance test (function `varImp` with default parameters in R package `caret`) was performed on the trained model to acquire the individual importance scores. Then antibodies were ranked by the importance scores, and 4 panels were used for antibody dropping test: (1) full 228 antibody panel; (2) top 100 most important antibodies; (3) top 50 most important antibodies; (4) top 30 most important antibodies.

#### CITE-seq BMC dataset

The CITE-seq healthy human BMC data with antibody panel of 25 markers was retrieved from R package `SeuratData` ‘bmcite’, orignated from Hao et al. (34). For benchmarking purpose, a total of 20k cells were randomly sampled from the original dataset, and selected for benchmarking. The first 15 components of the embedding vectors produced by all methods were used for benchmarking metric calculation. The UMAP visualization of the integration process was also calculated with the first 15 components of the embedding vectors. The original cell type annotation (lv2) from the R package was binned into 8 populations: “DC”, “progenitor”, “monocyte”, “NK”, “B”, “CD4 T”, “CD8 T” and “Other T”, and used for benchmarking.

#### Ab-seq BMC dataset

The Ab-seq healthy human BMC data with antibody panel of 97 markers, and whole transcriptome sequencing was retrieved from Triana et al. (39). All cells in the dataset (∼ 13k), except cells belonging to cell types with insufficient amount of cells (< 50 cells, annotated as “Doublet and Triplets”, “Early GMP”, “Gamma delta T cells”, “Immature B cells”, “Metaphase MPPs”, “Neutrophils” in Triana et al.) were excluded for integration, and the remaining 14 cell types were used during benchmarking. The first 15 components of the embedding vectors produced by all methods were used for benchmarking metric calculation. The UMAP visualization of the integration process was also calculated with the first 15 components of the embedding vectors.

#### TEA-seq PBMC dataset

The TEA-seq neutrophil-depleted human PBMC dataset was retrieved from Swanson et al. (41) (GSM4949911). This dataset is stained with 46 antibodies and contains chromatin accessibility information. Cell type annotation was performed using R package `Seurat`(v4) WNN-multi-modal clustering pipeline: function `FindMultiModalNeighbors` was run on ADT PCA (first 25 components) and ATAC LSI (2-50 components, calculated by R package `Archr` (42)). Subsequently, function `FindClusters` was used to generate unsupervised clustering (with parameter `algorithm = 3,` total of 8 populations were identified (“Naive CD4”, “Mem CD4”, “Monocyte”, “NK”, “Naive CD8”, “Mem CD8”, “Effector CD8”, “B”, “NK”), and the total amount of cells was ∼7.4k. ADT expressions and gene activity scores (calculated by R package `Archr` (42)) were used as input for MaxFuse and other methods. Additionally, during matching refinement, MaxFuse used LSI reduction of the ATAC peaks (first 2-50 components) as features for the ATAC modality. The first 15 components of the embedding vectors produced by all methods were used for benchmarking metric calculation. The UMAP visualization of the integration process was also calculated with the first 15 components of the embedding vectors.

### ASAP-seq PBMC dataset

The ASAP-seq healthy human PBMC data (CD28 & CD3 stim PBMC control group) with an antibody panel of 227 markers, and chromatin accessibility information was retrieved from Mimitou et al. (40) (GSM4732109). Cell type annotation was performed using R package `Seurat`(v4) WNN-multi-modal clustering pipeline: function `FindMultiModalNeighbors` was run on ADT PCA (first 18 components) and ATAC LSI (2-40 components, calculated by R package `Archr`). Subsequently, function `FindClusters` was used to generate unsupervised clustering (with parameter `algorithm = 3,`total of 9 populations were identified (“Naive CD4”, “Mem CD4”, “Monocyte”, “NK”, “Naive CD8”, “Mem CD8”, “B”, “OtherT”, “dirt”), and “dirt” was removed from subsequent usage, resulting in ∼ 4.4 k cells used. ADT expressions and gene activity scores (calculated by R package `Archr`) were used as input for MaxFuse and other methods. Additionally, during matching refinement, MaxFuse used LSI reduction of the ATAC peaks (First 2-50 components) as features for the ATAC modality. The first 15 components of the embedding vectors produced by all methods were used for benchmarking metric calculation. The UMAP visualization of the integration process was also calculated with the first 15 components of the embedding vectors.

### MaxFuse on Spatial-omics matching

#### CODEX and scRNA-seq human tonsil

CODEX multi - plexed imaging data of human tonsil tissues with a panel of 46 antibodies were retrieved from Kennedy-Darling et al. (47). Images from tonsil-9338 (region X2-8, Y7-15) were used. Whole-cell segmentation was performed with a local implementation of Mesmer (61), with weights downloaded from: https://deepcell-data.s3-us-west-1.amazonaws.com/model-weig'hts/Multiplex_Segmentation_20200908_2_head.h5. Inputs of segmentation were DAPI (nuclear) and CD45 (membrane). Signals from the images were capped at 99.7th percentile, with prediction parameter `model_mpp = 0.8`. Cells smaller than 30 pixels or larger than 800 pixels were excluded. Signals from individual cells were then extracted, and scaled to the [0, 1] interval, with percentile cutoffs of 0.5% (floor) and 99.5% (ceiling). Cell type annotation was performed using R package `Seurat` clustering pipeline: function `FindNeighbors` was run on CODEX protein PCA (first 15 components). Subsequently, function `FindClusters` was used to generate unsupervised clustering (with parameter `resolution = 1`), followed by manual annotation. A total of 9 populations were identified (“B-CD22-CD40”, “B-Ki67”, “Plasma”, “CD4 T”, “CD8 T”, “DC”, “Fibro/Epi”, “Vessel”, “Other”, and “Dirt”), and 6 populations (∼180k cells) were used in subsequent analysis (“B-CD22-CD40”, “B-Ki67”, “Plasma”, “CD4 T”, “CD8 T”, and “DC”).

Single-cell RNA-seq data of dissociated human tonsil cells were retrieved from King et al. (48). The pre-processing and cell typing steps were done in R package `Seurat`, following the description presented in King et al. In brief, tonsil cells (“t1”, “t2” and “t3”) were merged, then filtered by criteria: `nFeature_RNA > 200 & nFeature_RNA < 7500 & percent.mt < 20`, and subsequently value normalized by function `SCTransform`. Harmony batch correction was performed for different tonsils, with function `RunHarmony`. Unsupervised clustering was performed by function `FindNeighbors` with harmony embedding (1-27 dimensions) and function `FindClusters` with `resolution = 0.5`. A total of 8 population was defined (“B-CD22-CD40”, “B-Ki67”, “circulating B”, “Plasma”, “CD4 T”, “CD8 T”, “DC”, “Other”), and 6 populations (∼13k cells) were used in subsequent analysis (“B-CD22-CD40”, “B-Ki67”, “Plasma”, “CD4 T”, “CD8 T”, and “DC”).

Boundaries of germinal centers from the CODEX images were drawn manually, and dilation and erosion from the boundary was performed with python package `skimage`, with function `morphology.binary_dilation` and `morphology.disk`. Ten layers inward or outward from the boundary (each layer = 30 pixels, resolution: 376*nm*/pixel) was performed. Cells were assigned to each layer by their centroids’ locations. The RNA expression level from each layer, based on the averaged CODEX matched scRNA-seq cells, were plotted with R package `ggplot2`. The UMAP visualization of the integration process was calculated with the first 15 components of the embedding vectors.

#### HUBMAP atlas: CODEX, snRNA-seq and snATAC-seq human intestine

CODEX multiplex imaging (48 markers), snRNA-seq and snATAC-seq of healthy human intestine cells were acquired from Hickey et al. (32). For CODEX, samples “B005_SB” and “B006_CL” were used, while for snRNA-seq and snATAC-seq, single-ome sequencing data of four donors (“B001”, “B004”, “B005”, “B006”) from the study were used. Cells annotated as “B cells”, “T cells”, “Endothelial”, “Enteroendocrine”, “Goblet”, “Mono_Macrophages”, “Plasma”, “Smooth muscle”, and “Stroma” were selected for the integration process. Cell counts for each modality used for MaxFuse were: CODEX ∼100k (small bowel) and ∼70k (colon); snRNA-seq ∼32k (small bowel) and ∼16k (colon); snATAC-seq ∼28k (small bowel) and ∼21k (colon). CODEX protein expressions, snRNA-seq RNA expressions, snATAC-seq gene activity scores and LSI scores (calculated with R package `Archr`) were used as MaxFuse input (RNA expressions, gene activity scores and LSI scores were batch-corrected by Harmony (20), based on patient ID). The matching and integration process was done on colon and small bowel samples respectively.

Pairwise MaxFuse alignments of cells between protein (CODEX) and RNA (snRNA-seq), and cells between RNA (snRNA-seq) and ATAC (snATAC-seq) were performed. Relined pivots from the two bi-modal alignments were chained together by using the pivot cells in the RNA modality as the intermediary, resulting in a list of tri-modal pivots linking all three modalities. Subsequently, we used these pivots to calculate a tri-omic embedding via generalized CCA (gcca) (21, 55). In particular, we used the gcca formulation and algorithm described in (21).

The UMAP visualization of the tri-modal integration was calculated with the first 15 components of the embedding vectors (gcca scores in this case). Embeddings of CODEX cells were overlaid with their protein expressions, or their matched cells’ RNA expressions, or gene activity scores. Spatial locations of these expression values and scores were plotted based on CODEX cells’ x-y centroid locations. Additionally, we showed spatial locations of transcription factor motif enrichment scores (Z-score) of CODEX cells, based on their matched snRNA-seq cells, which were calculated by R package `chromVAR` (56). All values were capped between 5% — 95% quantiles for visualization purpose during plotting.

### Benchmark on ground-truth strongly linked modalities

#### MaxFuse and other methods specialized in ATAC-RNA integration in comparison

We compared MaxFuse to three methods that specialize in ATAC-RNA integration: scGLUE (19), Maestro (62) and scJoint (63). For MaxFuse, the initial matching used the gene activity scores, while during refined matching the active RNA features and LSI embedding from ATAC were used. For other methods in comparison, we used their default settings. Metrics used for benchmarking were calculated similarly as described in previous sections. The full detail (including preprocessing, implementation, and downstream analysis and evaluation of MaxFuse and other methods specialized in ATAC-RNA integration) is recorded and can be reproduced.

#### Multiome scRNA - scATAC-seq human retina dataset

Multiome (scRNA-seq & scATAC-seq) data of human retina cells was retrieved from Wang et al. (46). For input required by MaxFuse: gene activity and LSI scores of ATAC cells were calculated by R package `Archr` using the fragment files, while RNA counts were directy extracted. For other methods in comparison, we used their default settings. For benchmarking, a total of 20k cells were randomly sampled and used for testing. All cell types were used during integration (“Rod”, “OFF cone bipolar”, “Mullerglia”, “ON cone bipolar”, “Rod bipolar”, “Cone”, “GABA amacrine”, “Horizontal”, “Glyamacrine”, “AII amacrine”, “Retinal ganglion cell”, “Astrocyte”, “Microglia”, annotated by Wang et al.). The first 15 components of the embedding vectors produced by all methods were used for benchmarking metric calculation.

#### 10x Multiome peripheral blood mononuclear cells

Multiome (scRNA-seq & scATAC-seq) data of human mononuclear peripheral blood cells was retrieved from the 10x public data repository (44). For input required by MaxFuse: gene activity and LSI scores of ATAC modality were calculated by R package `Signac`, the latter using the fragment files. RNA counts were directly extracted from the `cellranger` output. Cell-eq CITE-seq PBMC reference (34) using the method in (34).

#### 10x Multiome day 18 embryonic mouse brain cells

Multiome (scRNA-seq & scATAC-seq) data of developing mouse brain cells was retrieved from the 10x public data repository (44). For input required by MaxFuse: gene activity and LSI scores of ATAC modality were calculated by R package `Signac`, the latter using the fragment files. RNA counts were directly extracted from the `cellranger` output. Cell-type labels were transferred from (64) using the method in (65).

#### 10x Multiome developing human cerebral cortex cells

Multiome (scRNA-seq & scATAC-seq) data of developing human cerebral cortex cells was retrieved from Trevino et al. (45). For input required by MaxFuse: gene activity and LSI scores of ATAC modality were calculated by R package `Signac` using the fragment files. RNA counts and ATAC peak matrices were extracted from 10x `cellranger` output. The cell-type labels were taken from the original publication.

## ACKNOWLEDGEMENTS

We thank Yuchao Jiang for initial discussions and for sharing the pre-processed data for 10x multiome PBMC and embryonic mouse brain datasets. B.Z. is supported by a Stanford Graduate Fellowship. This work was funded in part by grants from the National Science Foundation DMS-2210104 (Z.M.), the National Institutes of Health R01-HG006137-11, U2C-CA233285 (N.R.Z.), Mark Foundation Center for Radiobiology and Immunology (N.R.Z.), the US Food and Drug Administration Medical Countermeasures Initiative contracts HHSF223201610018C and 75F40120C00176 (G.P.N.), the Parker Institute for Cancer Immunotherapy (G.P.N.), and the Rachford and Carlota A. Harris Endowed Professorship (G.P.N.). This article reflects the views of the authors and should not be construed as representing the views or policies of the NSF, FDA, NIH, BMGF, Botnar Foundation or other institutions who provided funding.

## AUTHOR CONTRIBUTIONS

Conceptualization: S.C., B.Z., G.P.N., N.R.Z., Z.M.

Algorithm Development and Implementation: S.C., N.R.Z., Z.M.

Analysis: S.C., B.Z., S.H., Z.M.

Contribution of Key Reagents and Tools: J.W.H., K.Z.L., M.S., W.J.G, G.P.N.

Supervision: G.P.N., N.R.Z., Z.M.

Both S.C. and B.Z. contributed equally and have the right to list their name first in their CV.

## Reference

1. Marlon Stoeckius, Christoph Hafemeister, William Stephenson, Brian Houck-Loomis, Pratip K Chattopadhyay, Harold Swerdlow, Rahul Satija, and Peter Smibert. Simultaneous epitope and transcriptome measurement in single cells. Nature methods, 14(9):865–868, 2017.

2. Payam Shahi, Samuel C Kim, John R Haliburton, Zev J Gartner, and Adam R Abate. Ab-seq: Ultrahigh-throughput single cell protein profiling with droplet microfluidic barcoding. Scientific reports, 7(1):1–12, 2017.

3. Dominic Grün and Alexander van Oudenaarden. Design and analysis of single-cell sequencing experiments. Cell, 163(4):799–810, 2015.

4. Ino D Karemaker and Michiel Vermeulen. Single-cell dna methylation profiling: technologies and biological applications. Trends in biotechnology, 36(9):952–965, 2018.

5. Marek Bartosovic, Mukund Kabbe, and Gonçalo Castelo-Branco. Single-cell cut&tag profiles histone modifications and transcription factors in complex tissues. Nature biotechnology, 39(7):825–835, 2021.

6. Sebastian Preissl, Kyle J Gaulton, and Bing Ren. Characterizing cis-regulatory elements using single-cell epigenomics. Nature Reviews Genetics, pages 1–23, 2022.

7. Wai Lim Ku, Kosuke Nakamura, Weiwu Gao, Kairong Cui, Gangqing Hu, Qingsong Tang, Bing Ni, and Keji Zhao. Single-cell chromatin immunocleavage sequencing (scchic-seq) to profile histone modification. Nature methods, 16(4):323–325, 2019.

8. Caleb A Lareau, Fabiana M Duarte, Jennifer G Chew, Vinay K Kartha, Zach D Burkett, Andrew S Kohlway, Dmitry Pokholok, Martin J Aryee, Frank J Steemers, Ronald Lebofsky, et al. Droplet-based combinatorial indexing for massive-scale single-cell chromatin accessibility. Nature Biotechnology, 37(8):916–924, 2019.

9. Anjali Rao, Dalia Barkley, Gustavo S França, and Itai Yanai. Exploring tissue architecture using spatial transcriptomics. Nature, 596(7871):211–220, 2021.

10. Yury Goltsev, Nikolay Samusik, Julia Kennedy-Darling, Salil Bhate, Matthew Hale, Gustavo Vazquez, Sarah Black, and Garry P Nolan. Deep profiling of mouse splenic architecture with codex multiplexed imaging. Cell, 174(4):968–981, 2018.

11. Michael Angelo, Sean C Bendall, Rachel Finck, Matthew B Hale, Chuck Hitzman, Alexander D Borowsky, Richard M Levenson, John B Lowe, Scot D Liu, Shuchun Zhao, et al. Multiplexed ion beam imaging of human breast tumors. Nature medicine, 20(4):436–442, 2014.

12. Charlotte Giesen, Hao AO Wang, Denis Schapiro, Nevena Zivanovic, Andrea Jacobs, Bodo Hattendorf, Peter J Schüffler, Daniel Grolimund, Joachim M Buhmann, Simone Brandt, et al. Highly multiplexed imaging of tumor tissues with subcellular resolution by mass cytometry. Nature methods, 11(4):417–422, 2014.

13. Shanshan He, Ruchir Bhatt, Carl Brown, Emily A Brown, Derek L Buhr, Kan Chantranuvatana, Patrick Danaher, Dwayne Dunaway, Ryan G Garrison, Gary Geiss, et al. High-plex imaging of rna and proteins at subcellular resolution in fixed tissue by spatial molecular imaging. Nature Biotechnology, pages 1–13, 2022.

14. Emma Lundberg and Georg HH Borner. Spatial proteomics: a powerful discovery tool for cell biology. Nature Reviews Molecular Cell Biology, 20(5):285–302, 2019.

15. Yanxiang Deng, Marek Bartosovic, Sai Ma, D. Zhang, Petra Kukanja, Yang Xiao, Graham Su, Yang Liu, Xiaoyu Qin, Gorazd B Rosoklija, et al. Spatial profiling of chromatin accessibility in mouse and human tissues. Nature, 609(7926):375–383, 2022.

16. Ricard Argelaguet, Anna SE Cuomo, Oliver Stegle, and John C Marioni. Computational principles and challenges in single-cell data integration. Nature biotechnology, 39(10): 1202–1215, 2021.

17. Yang Xu and Rachel Patton McCord. Diagonal integration of multimodal single-cell data: potential pitfalls and paths forward. Nature Communications, 13(1):1–4, 2022.

18. Jinzhuang Dou, Shaoheng Liang, Vakul Mohanty, Xuesen Cheng, Sangbae Kim, Jongsu Choi, Yumei Li, Katayoun Rezvani, Rui Chen, and Ken Chen. Unbiased integration of single cell multi-omics data. BioRxiv, 2020.

19. Zhi-Jie Cao and Ge Gao. Multi-omics single-cell data integration and regulatory inference with graph-linked embedding. Nature Biotechnology, pages 1–9, 2022.

20. Ilya Korsunsky, Nghia Millard, Jean Fan, Kamil Slowikowski, Fan Zhang, Kevin Wei, Yuriy Baglaenko, Michael Brenner, Po-ru Loh, and Soumya Raychaudhuri. Fast, sensitive and accurate integration of single-cell data with harmony. Nature methods, 16(12):1289–1296, 2019.

21. Bokai Zhu, Shuxiao Chen, Yunhao Bai, Han Chen, Nilanjan Mukherjee, Gustavo Vazquez, David R McIlwain, Alexandar Tzankov, Ivan T Lee, Matthias S Matter, et al. Robust single-cell matching and multi-modal analysis using shared and distinct features reveals orchestrated immune responses. bioRxiv, 2021.

22. Joshua D Welch, Velina Kozareva, Ashley Ferreira, Charles Vanderburg, Carly Martin, and Evan Z Macosko. Single-cell multi-omic integration compares and contrasts features of brain cell identity. Cell, 177(7):1873–1887, 2019.

23. Kevin E Wu, Kathryn E Yost, Howard Y Chang, and James Zou. Babel enables cross-modality translation between multiomic profiles at single-cell resolution. Proceedings of the National Academy of Sciences, 118(15):e2023070118, 2021.

24. Tim Stuart, Andrew Butler, Paul Hoffman, Christoph Hafemeister, Efthymia Papalexi, William M Mauck III, Yuhan Hao, Marlon Stoeckius, Peter Smibert, and Rahul Satija. Comprehensive integration of single-cell data. Cell, 177(7):1888–1902, 2019.

25. Zhen Miao, Benjamin D Humphreys, Andrew P McMahon, and Junhyong Kim. Multi-omics integration in the age of million single-cell data. Nature Reviews Nephrology, 17(11):710– 724, 2021.

26. Jinzhuang Dou, Shaoheng Liang, Vakul Mohanty, Qi Miao, Yuefan Huang, Qingnan Liang, Xuesen Cheng, Sangbae Kim, Jongsu Choi, Yumei Li, et al. Bi-order multimodal integration of single-cell data. Genome biology, 23(1):1–25, 2022.

27. Zhana Duren, Xi Chen, Mahdi Zamanighomi, Wanwen Zeng, Ansuman T Satpathy, Howard Y Chang, Yong Wang, and Wing Hung Wong. Integrative analysis of single-cell genomics data by coupled nonnegative matrix factorizations. Proceedings of the National Academy of Sciences, 115(30):7723–7728, 2018.

28. Vivien Marx. A dream of single-cell proteomics. Nature Methods, 16(9):809–812, 2019.

29. Vidhya M. Ravi, Paulina Will, Jan Kueckelhaus, Na Sun, Kevin Joseph, Henrike Salié, Lea Vollmer, Ugne Kuliesiute, Jasmin von Ehr, Jasim K. Benotmane, Nicolas Neidert, Marie Follo, Florian Scherer, Jonathan M. Goeldner, Simon P. Behringer, Pamela Franco, Mo-hammed Khiat, Junyi Zhang, Ulrich G. Hofmann, Christian Fung, Franz L. Ricklefs, Ka-trin Lamszus, Melanie Boerries, Manching Ku, Jürgen Beck, Roman Sankowski, Marius Schwabenland, Marco Prinz, Ulrich Schüller, Saskia Killmer, Bertram Bengsch, Axel K. Walch, Daniel Delev, Oliver Schnell, and Dieter Henrik Heiland. Spatially resolved multi-omics deciphers bidirectional tumor-host interdependence in glioblastoma. Cancer Cell, 40(6):639–655.e13, 2022. ISSN 1535-6108. doi: https://doi.org/10.1016/j.ccell.2022.05.009.

30. Amin Abedini, Ziyuan Ma, Julia Frederick, Poonam Dhillon, Michael S Balzer, Rojesh Shrestha, Hongbo Liu, Steven Vitale, Kishor Devalaraja-Narashimha, Paola Grandi, et al. Spatially resolved human kidney multi-omics single cell atlas highlights the key role of thefibrotic microenvironment in kidney disease progression. bioRxiv, 2022.

31. Anuja Sathe, Kaishu Mason, Susan M Grimes, Zilu Zhou, Billy T Lau, Xiangqi Bai, Andrew Su, Xiao Tan, H Lee, Carlos J Suarez, et al. Colorectal cancer metastases in the liver estab-lish immunosuppressive spatial networking between tumor associated spp1+ macrophagesand fibroblasts. Clinical Cancer Research: an Official Journal of the American Association for Cancer Research, pages CCR–22, 2022.

32. John W Hickey, Winston R Becker, Stephanie A Nevins, Aaron Horning, Almudena EspinPerez, Roxanne Chiu, Derek C Chen, Daniel Cotter, Edward D Esplin, Annika K Weimer, et al. High resolution single cell maps reveals distinct cell organization and function acrossdifferent regions of the human intestine. bioRxiv, 2021.

33. Rainer Burkard, Mauro Dell’Amico, and Silvano Martello. Assignment problems: revised reprint. SIAM, 2012.

34. Yuhan Hao, Stephanie Hao, Erica Andersen-Nissen, William M Mauck III, Shiwei Zheng,Andrew Butler, Maddie J Lee, Aaron J Wilk, Charlotte Darby, Michael Zager, et al. Integratedanalysis of multimodal single-cell data. Cell, 184(13):3573–3587, 2021.

35. Hoa Thi Nhu Tran, Kok Siong Ang, Marion Chevrier, Xiaomeng Zhang, Nicole Yee ShinLee, Michelle Goh, and Jinmiao Chen. A benchmark of batch-effect correction methods forsingle-cell rna sequencing data. Genome biology, 21(1):1–32, 2020.

36. Jie Liu, Yuanhao Huang, Ritambhara Singh, Jean-Philippe Vert, and William Stafford Noble. Jointly embedding multiple single-cell omics measurements. In Algorithms in bioinformat ics:… International Workshop, WABI…, proceedings. WABI (Workshop), volume 143. NIHPublic Access, 2019.

37. April R Kriebel and Joshua D Welch. Uinmf performs mosaic integration of single-cell multi-omic datasets using nonnegative matrix factorization. Nature communications, 13(1):1–17, 2022.

38. Etienne Becht, Leland McInnes, John Healy, Charles-Antoine Dutertre, Immanuel WHKwok, Lai Guan Ng, Florent Ginhoux, and Evan W Newell. Dimensionality reduction forvisualizing single-cell data using umap. Nature biotechnology, 37(1):38–44, 2019.

39. Sergio Triana, Dominik Vonficht, Lea Jopp-Saile, Simon Raffel, Raphael Lutz, Daniel Leonce, Magdalena Antes, Pablo Hernández-Malmierca, Diana Ordoñez-Rueda, Beáta Ra-masz, et al. Single-cell proteo-genomic reference maps of the hematopoietic system enablethe purification and massive profiling of precisely defined cell states. Nature immunology,22(12):1577–1589, 2021.

40. Eleni P Mimitou, Caleb A Lareau, Kelvin Y Chen, Andre L Zorzetto-Fernandes, Yuhan Hao,Yusuke Takeshima, Wendy Luo, Tse-Shun Huang, Bertrand Z Yeung, Efthymia Papalexi, et al. Scalable, multimodal profiling of chromatin accessibility, gene expression and proteinlevels in single cells. Nature biotechnology, 39(10):1246–1258, 2021.

41. Elliott Swanson, Cara Lord, Julian Reading, Alexander T Heubeck, Palak C Genge, Zachary Thomson, Morgan DA Weiss, Xiao-jun Li, Adam K Savage, Richard R Green, et al. Simulta-neous trimodal single-cell measurement of transcripts, epitopes, and chromatin accessibilityusing tea-seq. Elife, 10:e63632, 2021.

42. Jeffrey M Granja, M Ryan Corces, Sarah E Pierce, S Tansu Bagdatli, Hani Choudhry,Howard Y Chang, and William J Greenleaf. Archr is a scalable software package for inte-grative single-cell chromatin accessibility analysis. Nature genetics, 53(3):403–411, 2021.

43. Kevin Z. Lin and Nancy R. Zhang. Quantifying common and distinct information in single-cellmultimodal data with tilted-cca. bioRxiv, 2022. doi: 10.1101/2022.10.07.511320.

44. 10X Genomics. 10x genomics datasets, 2022.

45. Alexandro E Trevino, Fabian Müller, Jimena Andersen, Laksshman Sundaram, Arwa Kathiria, Anna Shcherbina, Kyle Farh, Howard Y Chang, Anca M Pas, ca, Anshul Kundaje, et al. Chromatin and gene-regulatory dynamics of the developing human cerebral cortex atsingle-cell resolution. Cell, 184(19):5053–5069, 2021.

46. Sean K Wang, Surag Nair, Rui Li, Katerina Kraft, Anusri Pampari, Aman Patel, Joyce BKang, Christy Luong, Anshul Kundaje, and Howard Y Chang. Single-cell multiome of thehuman retina and deep learning nominate causal variants in complex eye diseases. bioRxiv, 2022.

47. Julia Kennedy-Darling, Salil S Bhate, John W Hickey, Sarah Black, Graham L Barlow,Gustavo Vazquez, Vishal G Venkataraaman, Nikolay Samusik, Yury Goltsev, Christian MSchürch, et al. Highly multiplexed tissue imaging using repeated oligonucleotide exchangereaction. European Journal of Immunology, 51(5):1262–1277, 2021.

48. Hamish W King, Kristen L Wells, Zohar Shipony, Arwa S Kathiria, Lisa E Wagar, Caleb Lareau, Nara Orban, Robson Capasso, Mark M Davis, Lars M Steinmetz, et al. Integratedsingle-cell transcriptomics and epigenomics reveals strong germinal center–associated eti-ology of autoimmune risk loci. Science Immunology, 6(64):eabh3768, 2021.

49. Stella Maris Ranuncolo, Jose M Polo, Jamil Dierov, Michael Singer, Tracy Kuo, John Greally,Roland Green, Martin Carroll, and Ari Melnick. Bcl-6 mediates the germinal center b cell phenotype and lymphomagenesis through transcriptional repression of the dna-damagesensor atr. Nature immunology, 8(7):705–714, 2007.

50. Masayuki Kuraoka, T Matt Holl, Dongmei Liao, Mandy Womble, Derek W Cain, Alexander EReynolds, and Garnett Kelsoe. Activation-induced cytidine deaminase mediates centraltolerance in b cells. Proceedings of the National Academy of Sciences, 108(28):11560–11565, 2011.

51. Antony B Holmes, Clarissa Corinaldesi, Qiong Shen, Rahul Kumar, Nicolo Compagno, Zhong Wang, Mor Nitzan, Eli Grunstein, Laura Pasqualucci, Riccardo Dalla-Favera, et al. Single-cell analysis of germinal-center b cells informs on lymphoma cell of origin and out-come. Journal of Experimental Medicine, 217(10), 2020.

52. Dan Suan, Nike J Kräutler, Jesper LV Maag, Danyal Butt, Katherine Bourne, Jana R Hermes, Danielle T Avery, Clara Young, Aaron Statham, Michael Elliott, et al. Ccr6 defines memory b cell precursors in mouse and human germinal centers, revealing light-zone location and predominant low antigen affinity. Immunity, 47(6):1142–1153, 2017.

53. Sean P Saunders, Erica GM Ma, Carlos J Aranda, and Maria A Curotto de Lafaille. Non-classical b cell memory of allergic ige responses. Frontiers in immunology, 10:715, 2019.

54. Lyssia Belarif, Caroline Mary, Lola Jacquemont, Hoa Le Mai, Richard Danger, Jeremy Hervouet, David Minault, Virginie Thepenier, Veronique Nerrière-Daguin, Elisabeth Nguyen, et al. Il-7 receptor blockade blunts antigen-specific memory t cell responses and chronic inflammation in primates. Nature communications, 9(1):1–13, 2018.

55. Jon R Kettenring. Canonical analysis of several sets of variables. Biometrika, 58(3):433– 451, 1971.

56. Alicia N Schep, Beijing Wu, Jason D Buenrostro, and William J Greenleaf. chromVAR: inferring transcription-factor-associated accessibility from single-cell epigenomic data. Nature methods, 14(10):975–978, 2017.

57. Sorim Nam and Jong-Seok Lim. Essential role of interferon regulatory factor 4 (irf4) in immune cell development. Archives of pharmacal research, 39(11):1548–1555, 2016.

58. Jonathan P Katz, Nathalie Perreault, Bree G Goldstein, Catherine S Lee, Patricia A Labosky, Vincent W Yang, and Klaus H Kaestner. The zinc-finger transcription factor klf4 is required for terminal differentiation of goblet cells in the colon. 2002.

59. Zhigao Wang, Da-Zhi Wang, GC Teg Pipes, and Eric N Olson. Myocardin is a master regulator of smooth muscle gene expression. Proceedings of the National Academy of Sciences, 100(12):7129–7134, 2003.

60. Shuxiao Chen, Sizun Jiang, Zongming Ma, Garry P Nolan, and Bokai Zhu. One-way matching of datasets with low rank signals. arXiv preprint 2204.13858, 2022.

61. Noah F Greenwald, Geneva Miller, Erick Moen, Alex Kong, Adam Kagel, Thomas Dougherty, Christine Camacho Fullaway, Brianna J McIntosh, Ke Xuan Leow, Morgan Sarah Schwartz, et al. Whole-cell segmentation of tissue images with human-level performance using large-scale data annotation and deep learning. Nature biotechnology, 40(4):555–565, 2022.

62. Chenfei Wang, Dongqing Sun, Xin Huang, Changxin Wan, Ziyi Li, Ya Han, Qian Qin, Jingyu Fan, Xintao Qiu, Yingtian Xie, et al. Integrative analyses of single-cell transcriptome and regulome using maestro. Genome biology, 21(1):1–28, 2020.

63. Yingxin Lin, Tung-Yu Wu, Sheng Wan, Jean YH Yang, Wing H Wong, and YX Wang. scjoint integrates atlas-scale single-cell rna-seq and atac-seq data with transfer learning. Nature Biotechnology, 40(5):703–710, 2022.

64. Gioele La Manno, Kimberly Siletti, Alessandro Furlan, Daniel Gyllborg, Elin Vinsland, Alejandro Mossi Albiach, Christoffer Mattsson Langseth, Irina Khven, Alex R Lederer, Lisa M Dratva, et al. Molecular architecture of the developing mouse brain. Nature, 596(7870): 92–96, 2021.

65. Mo Huang, Zhaojun Zhang, and Nancy R Zhang. Dimension reduction and denoising of single-cell rna sequencing data in the presence of observed confounding variables. bioRxiv, 2020.

